# Stimulus-Locked Responses on Human Upper Limb Muscles and Corrective Reaches are Preferentially Evoked by Low Spatial Frequencies

**DOI:** 10.1101/690354

**Authors:** Rebecca A. Kozak, Philipp Kreyenmeier, Chao Gu, Kevin Johnston, Brian D. Corneil

## Abstract

In situations requiring immediate action, humans can generate visually-guided responses at remarkably short latencies. Here, to better understand the visual attributes that best evoke such rapid responses, we recorded upper limb muscle activity while participants performed visually-guided reaches towards Gabor patches composed of differing spatial frequencies. We studied reaches initiated from a stable posture (experiment 1, a static condition), or during on-line reach corrections to an abruptly displaced target (experiment 2, a dynamic condition). In both experiments, we detail the latency and prevalence of stimulus-locked responses (SLRs), which are brief bursts of EMG activity that are time-locked to target presentation rather than movement onset. SLRs represent the first wave of EMG recruitment influenced by target presentation, and enable quantification of rapid visuomotor transformations. In both experiments, reach targets composed of low spatial frequencies elicited the shortest latency and most prevalent SLRs, with SLR latency increasing and SLR prevalence decreasing for reach targets composed of progressively higher spatial frequencies. SLRs could be evoked in either the static or dynamic condition, and when present in experiment 2, were associated with shorter latency and larger magnitude corrections. Furthermore, SLRs evolved at shorter latencies (~20 ms) when the arm was already in motion. These results demonstrate that stimuli composed of low spatial frequencies preferentially evoke the most rapid visuomotor responses which, in the context of rapidly correcting an on-going reaching movement, are associated with earlier and larger on-line reach corrections.

**Significance Statement:** Humans have a remarkable capacity to respond quickly to changes in our visual environment. Although our visual world is composed of a range of spatial frequencies, surprisingly little is known about which frequencies preferentially evoke rapid reaching responses. Here, we systematically varied the spatial frequency of peripheral reach targets while measuring EMG activity on an upper limb muscle. We found that visual stimuli composed of low-spatial frequencies elicit the most rapid and robust EMG responses and corrective reaches. Thus, when time is of the essence, low spatial frequencies preferentially drive fast visuomotor responses.

## Introduction

To reach towards a visible target, visual information is ultimately transformed into motor commands. Doing so requires extraction of visual attributes such as the color, shape, and size of a visible target, and integration of these features into a command that is relayed to the motor periphery typically via the corticospinal tract. However, there are instances where we have to respond as quickly as possible, for example, to catch a falling mug or to volley a tennis ball that deflects off the net during a tennis match. Which features of a visual stimulus best engender such rapid visuomotor responses?

One way to study this question in the laboratory is to examine the latencies at which participants respond to targets that are suddenly displaced during an on-going reaching movement. In such dynamic scenarios, humans can initiate on-line reach corrections within a remarkably short latency of ~125 ms following the target displacement (Soechting and Lacquaniti, 1983; Day and Lyon, 2000). Veerman and colleagues (2008) systematically investigated the influence of visual attributes in online reach corrections, and reported that earlier responses were driven by stimuli defined by luminance, orientation, and size, but not other attributes such as color. These results demonstrate that some visual attributes, but not others, are crucial for these rapid online reach corrections.

Recently, we have described a novel way of investigating rapid visuomotor responses which can be studied from a static posture. *Stimulus-locked responses* (SLRs) are short-latency bursts of directionally-tuned electromyographic (EMG) activity that evolve time-locked within ~100 ms of stimulus onset (Pruszynski et al., 2010), persisting even if an eventual reach is withheld (Wood et al., 2015; Atsma et al., 2018). Although never studied in the same paradigm, there are many similarities in the response properties of SLRs and on-line corrections. For example, both SLRs (Gu et al., 2016) and the trajectory of the initial phase of on-line corrections (Day and Lyon, 2000) are inexorably drawn toward the visual target, even when participants are instructed to move in the opposite direction. We and others have hypothesized that SLRs, and by extension the initial phase of an on-line reach corrections, are the product of a tecto-reticulospinal pathway that is mediated through the intermediate and deep layers of the superior colliculus (SCi) (Pruszynski et al., 2010; Corneil and Munoz, 2014). Consistent with this, SLRs (Wood et al., 2015), online reach corrections (Veerman et al., 2008), and visual responses within the SCi (Marino et al., 2012), are all evoked at a shorter latency by high contrast stimuli.

The current study was motivated by a recent paper which reported that visual responses in the SCi responses are dependent on the spatial frequency (SF) of a visual stimulus, with SCi neurons responding sooner to lower SFs (Chen et al., 2018). Our hypothesis that the visual response in the SCi drives both the SLR and the earliest component of on-line reach corrections predicts that both SLRs and on-line reach corrections should evolve sooner for lower SF stimuli. We test this prediction in two experimental frameworks, measuring SLRs alone in reaches starting form a static posture (*experiment 1*), and measuing both SLRs and on-line reach corrections in a dynamic task where a reach target is displaced in mid-flight (*experiment 2*). Consistent with our hypothesis, we find that rapid visuomotor responses, whether indexed by SLRs or the latency of on-line reach corrections, are preferentially evoked by stimuli composed of low SFs.

## Materials and methods

### Participants

A total of 33 participants (18 females, 15 males; mean age: 23.9 years SD: 3.4) completed at least one of two experiments. All participants provided informed consent, were paid for their participation, and were free to withdrawal from the experiment at any time. All but two participants were right-handed. All participants had normal or corrected-to-normal vision, with no current visual, neurological, or musculoskeletal disorders. All procedures were approved by the Health Science Research Ethics Board at the University of Western Ontario.

### Apparatus

Participants were seated in a KINARM robotic exoskeleton (BKIN Technologies, Kingston, ON, Canada), and performed reaching movements with their right arm (**Fig. 1a**). As previously described (Scott, 1999; Pruszynski et al., 2008), this robot permits arm movements in a horizontal plane via shoulder and elbow flexion and extension, and allows for torque application at these joints. Visual stimuli were projected from a downward facing LED monitor (LG 47LS35A, size: 47” resolution: 1920×1080 pixels), onto an upward facing mirror. A shield underneath the mirror occluded direct vision of the hand, but real-time hand position was represented on the monitor via a black dot with a diameter of 1 cm. A photodiode was used to indicate the precise time of onset of the peripheral visual stimulus, and all kinematic and electromyographic (EMG) data were aligned to photodiode onset. Eye movements were not measured.

**Figure 1.**
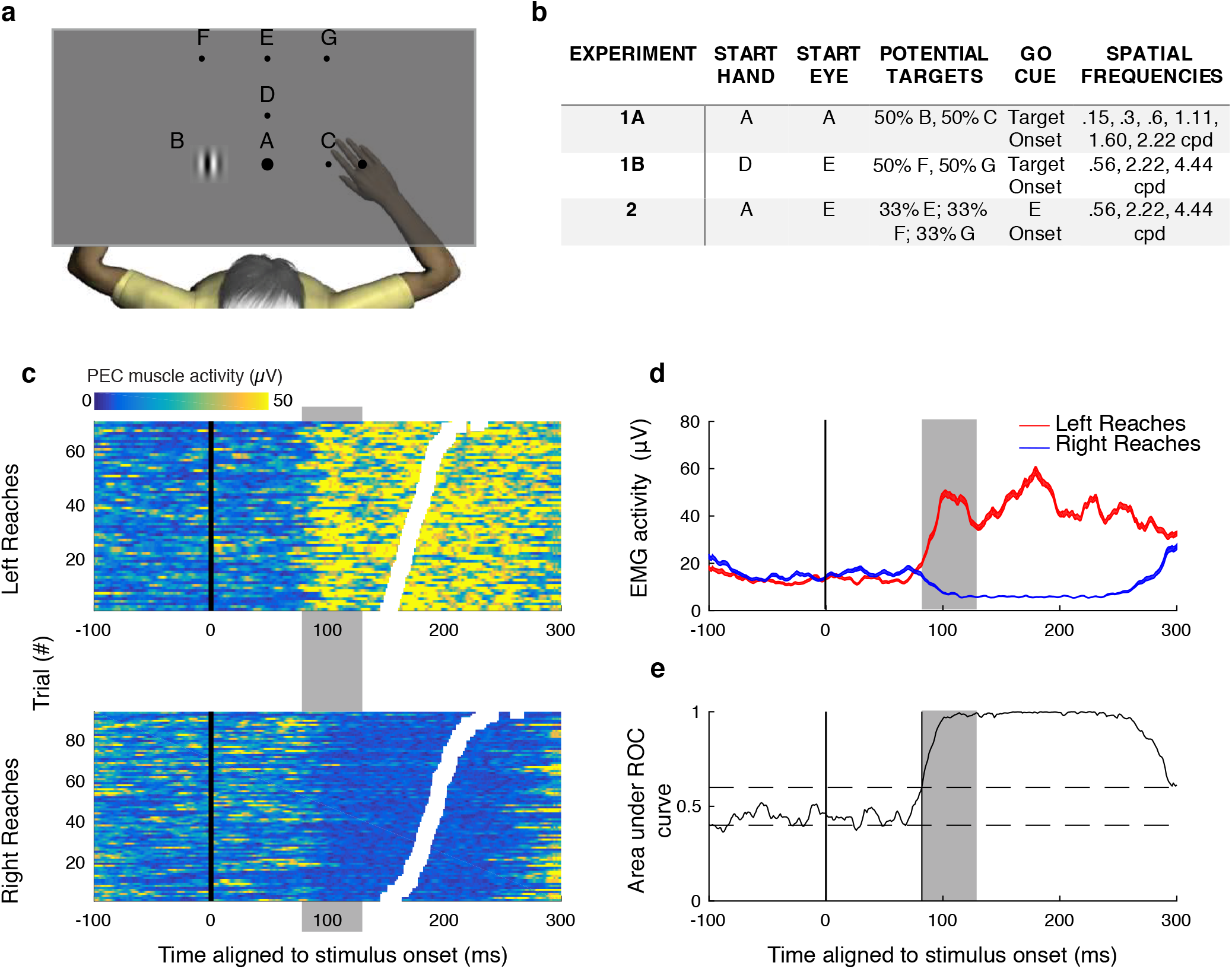
Experimental design and SLR classification. (a, b) General display and configuration of targets (a) and corresponding hand and eye start positions, potential targets, go cues and spatial frequencies used for each of the experiments (b; see Methods for more details). (c) Trial-by-trial recruitment of right PEC for an example subject generated left or right reaches. Each row is a different trial, with colour conveying degree of recruitment. Trials sorted by RT (white boxes), and aligned to stimulus onset. The SLR (highlighted in grey vertical box) appears as a vertical banding of increased or decreased recruitment aligned to left or right stimulus presentation, respectively, rather than movement onset. (d, e) Mean EMG activity (+/− standard error; d) and time-series ROC analysis (e) for data shown in (c). Horizontal dashed lines in e shows the 0.6 or 0.4 level which constitutes discrimination threshold.

### Experimental design

#### Experiment 1a: Static Task

Participants (*n*= 16) performed visually guided reaches (**Fig. 1a,b**) towards peripheral stimuli composed of stationary Gabor patches (see below), starting from a static posture in which the hand was not in motion. Throughout the experiment, a constant torque of 2 Nm was applied to the shoulder for the entire experiment, increasing tonic activity on the muscle of interest. Doing so allowed for observations in decreases of muscle recruitment when the arm moved in the non-preferred direction for the muscle of interest. Participants initiated each trial by bringing their hand into a central black stimulus, which was located 45ccm in front of them relative to their midline. This central stimulus had a 1 cm diameter and changed to white once the hand attained this position. Participants were instructed to foveate this start position before target onset. The trial was reset if the hand exited the start position before completion of a hold period (randomized from 1-1.5 s). Simultaneous with the disappearance of the start position stimulus, a Gabor patch subtending 7 cm then appeared either 10 cm to the left or right of the start position; the center of this patch was at an eccentricity of ~9° to the left or right of the start position; subtending approximately 12°. The Gabor patch was composed of one of six spatial frequencies (SF) ranging between 0.15 to 2.22 cycles per degree (cpd) (**Fig. 1b**; see below for details on how Gabor patches were constructed). Our lowest SF was implemented as a control target, as we presented a very low SF, effectively presented as a gaussian blurred black dot. Participants were instructed to reach towards the peripheral target as quickly as possible, with the trial ending when the virtual cursor made contact with the Gabor. There were 12 unique trial conditions (six SFs, each presented to the left and right). Participants completed five blocks of 240 trials each, with each block containing 20 repetitions of each unique trial condition presented pseudorandomly without replacement, yielding a total of 100 trials for each unique trial condition.

#### Experiment 1b: Static task with higher SF targets

In experiment 1b, peripheral targets were placed at an increased distance from the subject, permitting the presentation of higher spatial frequencies than in experiment 1a. Stimuli were placed at locations resembling those used in experiment 2 (see below), so that the location of the targets relative to the eye and hand were approximately the same in experiments 1b and 2. 16 participants, some of whom participated in experiment 1a, also participated in experiment 1b (**Fig. 1a,b**). 14 of 16 participants in experiment 1b also participated in experiment 2 (see **Fig. 2b** for participant details).

**Figure 2.**
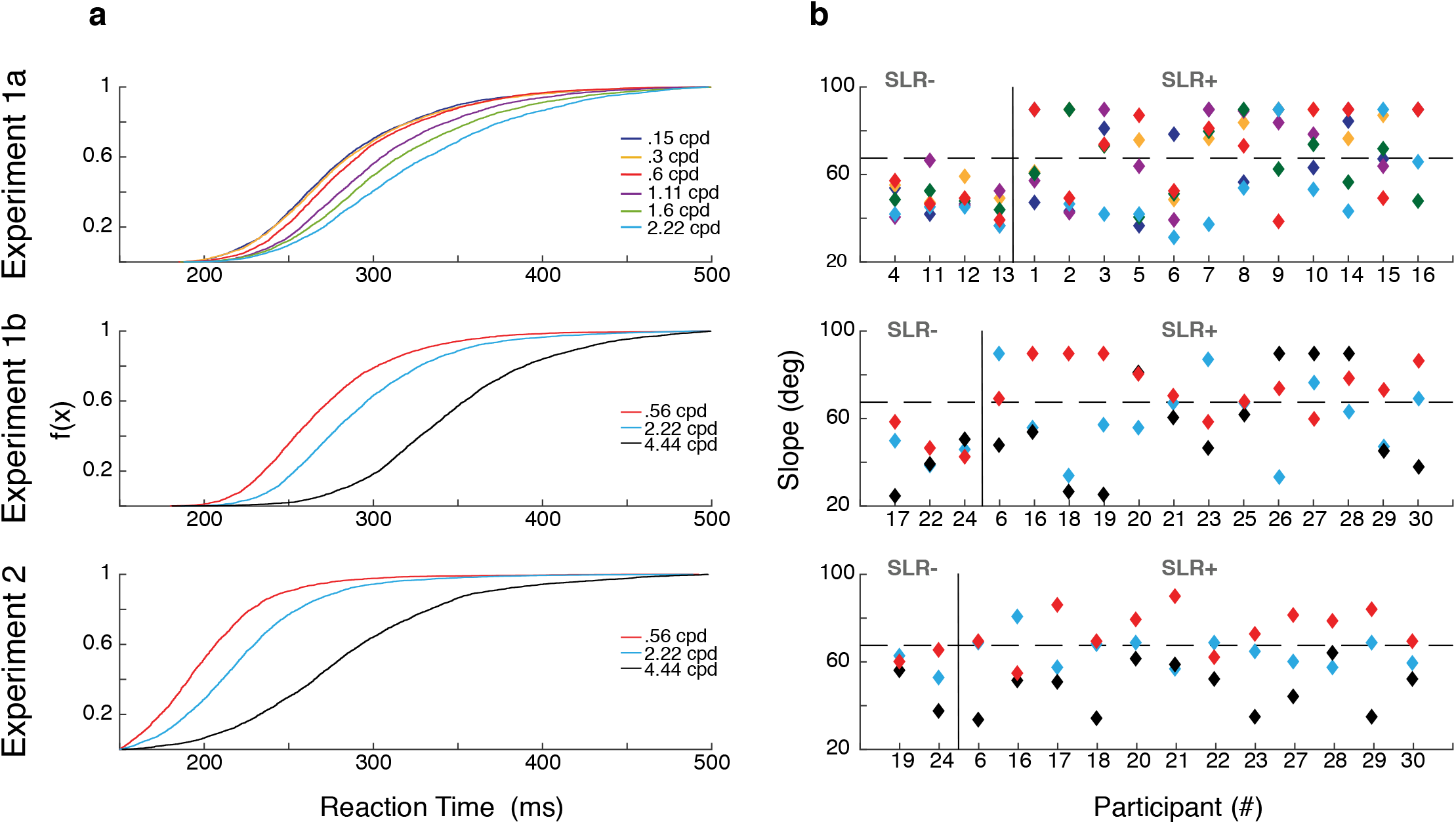
RTs and detection of SLRs. (a) Cumulative RT distributions in all experiments, pooled across all participants and directions and segregated by SF. (b) Slope of the relationship between discrimination time and average RT for early and late RT groups, for all experiments. Each data point represents a unique subject for a particular SF. Slopes are capped at 90 deg. Results are grouped into those who exhibiting an SLR in at least one condition (SLR+) or not (SLR−), given a slope threshold of 67.5 deg (horizontal dashed line).

Participants initiated each trial by bringing their hand into a start position (2 cm diameter) located 55.5 cm in front of them. A constant torque of 3Nm was applied to the shoulder for the entire experiment. A higher torque was used than in experiment 1a, as target positions were less in the preferred direction of the muscle. A fixation cross then appeared 10 cm above the central hold marker, which participants were instructed to look at while not moving their hand. Participants were required to maintain the initial hand position for a randomized period of 1-2 seconds. After this, a 5 cm diameter Gabor patch composed of one of three frequencies (.56, 2.22, 4.44 cpd; **Fig. 1b**) subtending approximately 4.5° of visual angle, was presented. The center of this patch was ~8° of visual angle to the left or right of the start position. Participants were instructed to reach and look to this peripheral target as quickly as possible. Participants completed four blocks of 150 trials each, with each block composed of 25 trials of each unique trial condition (three SFs to the right and left), yielding a total of 100 repeats of each unique trial condition.

#### Experiment 2: Dynamic Task

In experiment 2, participants (n = 14, all of whom also completed experiment 1b in the same session) occasionally generated online corrections to stimuli that suddenly jumped to the left or right just after the start of a reaching movement. We term this a dynamic task, as the hand was in motion when the target stimulus was displaced, necessitating an online correction. Peripheral stimuli were presented at the same screen locations as in experiment 1b, and initial eye and hand positions were also dissociated. A constant torque was not applied during experiment 2, as participants in pilot experiments found this load to be too difficult to maintain for the duration of experiment 2.

Participants initiated each trial by first bringing their hand into a central hold position (38 cm in front of them), while they looked at a central fixation cross located 17.5 cm above the central hold stimulus (**Fig. 1a,b**). The central hold stimulus for the hand was placed closer to the participant in experiment 2 than in experiment 1b, as a longer reaching movement permitted more time for a mid-flight correction. Further, this configuration meant that the mid-flight hand position at peripheral target presentation closely resembled that used in experiment 1b. Participants then maintained this dissociated hand and eye position for 1-2s. After this, the central fixation cross changed to a Gaussian blurred black dot, cueing the participant to reach toward this central location. All trial types were identical up to this time. On 1/3^rd^ of all trials (termed control trials), no other event occurred and participants were simply required to reach towards the gaussian blurred black dot. On the remaining 2/3rds of all trials (termed jump trials), the blurred Gaussian black dot was replaced by a peripheral target identical to those used in experiment 1b. The presentation of this peripheral target occurred when the hand exited the central hold position (1 cm diameter), requiring participants to adjust an on-going reach movement to either the left or right. Participants were instructed to reach as quickly as possible to the fixation target on control trials, and to try to correct their reach movements as quickly as possible on jump trials, so that they could move through the bottom aspect of the peripheral Gabor patch. There was a total of seven unique trial combinations: a control trial, and six different jump trials (three SFs per direction). Participants completed four blocks of 225 trials each, with each block composed pseudorandomly of 75 control trials and 25 repeats of each unique jump trial condition; thus, there were a total of 300 control trials and 100 trials of each unique jump trial condition.

#### Composition of peripheral Gabor target patches

All stimuli were created in Matlab using the Psychophysics toolbox (Brainard, 1997; Pelli, 1997). Due to monitor resolution and the viewing distance of the targets, we were limited to an upper limit of 2.22 cpd in experiment 1a, and 4.44 cpd in experiment 1b and 2. Stimuli consisted of vertical sinewaves overlaid with a Gaussian window. In all experiments, peripheral target stimuli consisted of perceptually contrast matched Gabor patches varying in SF measured in cpd of visual angle. Our motivation to implement a contrast matching procedure was two-fold. First, we performed this procedure in an attempt to isolate the effects of SF from perceived contrast, as contrast influences SLR latency and magnitude (Wood et al., 2015). Second, perceptual contrast matching mitigates some of the increases in RT with higher SFs (Breitmeyer, 1975), which was important given that larger magnitude SLRs precede shorter RTs (Pruszynski et al., 2010; Gu et al., 2016). We used a modified contrast matching procedure with foveal stimuli as described in (Davidson, 1968). A double random interleaved procedure was used for each SF, on a linearized (gamma-corrected) screen (mean background luminance 42.78 cd/m2). Participants were presented a fixation point, followed by a flashed Gabor patch (200 ms), followed by a pause of 500 ms, followed by a second flashed Gabor patch (200 ms); participants were instructed to “indicate the stimulus with more contrast”, by button press. In each experiment, we used a single standard stimulus presented at a suprathreshold 85% contrast from the middle SF used in each experiment, with the comparison stimulus from each of the SFs being composed of a Gabor patch presented at a higher or lower contrast, as determined by a staircase procedure. Our rationale for using suprathreshold contrast is based on the relationship between SLR detectability and contrast (Wood et al., 2015). After a reversal (participant changes response from increasing to decreasing contrast or vice versa), the step size was decreased. Each staircase procedure required ten reversals, and the contrast for a matched stimulus was taken as the mean contrast of the last five reversals. All participants completed this procedure for all SFs used in a given experiment, yielding a unique set of perceptually contrast-matched Gabors across a range of SFs for each subject, which were then used in the experiment.

#### Data acquisition and analysis

As previously described (Gu et al., 2018), surface electromyographic (EMG) activity was recorded from the clavicular head of the right pectoralis muscle with double-differential surface electrodes (Delsys Inc. Bagnoli-8 system, Boston, MA USA). We placed two electrodes on each participant, and chose the recording exhibiting the higher level of background activity. EMG signals were sampled at 1,000 Hz, amplified by 1000, and rectified off-line. We excluded data from three participants (one from experiment 1a, two from experiment 1b, and four from experiment 2) due to an absence or very low level of recruitment for movements in the preferred direction of the muscle. We used the same muscle recordings in all but three participants between experiments 1b and 2.

All kinematic data were sampled at 1000 Hz by the Kinarm data acquisition system. Reaction time (RT) was calculated as time from peripheral target appearance (measured by the photodiode) to the initiation of the reaching movement. In all experiments, reach initiation was determined using the lateral velocity towards left and right targets, extrapolating a line drawn between the crossings of 25% and 75% of single trial peak velocities back to zero (as in (Veerman et al., 2008)).

Our previous work on upper limb muscle recruitment during visually-guided reaching has distinguished the small burst of EMG activity aligned to stimulus onset (i.e., the SLR) from the larger, later burst of EMG activity associated with onset of the reaching movement (Gu et al., 2016, 2018). The distinction between the SLR and the later movement-related burst becomes blurred for shorter-RT movements. Given that the goal of this manuscript is to isolate the effects of the spatial frequency of a peripheral stimulus on the SLR, we excluded trials below a particular RT (excluding trials with RTs < 185 ms in experiments 1a and 1b, and trials with RTs < 150 ms in experiment 2), since RTs of on-line reach corrections tend to be shorter than RTs initiated from a static posture. In all experiments, we also excluded trials with RTs > 500 ms, due to presumed inattentiveness. Finally, in all experiments, we excluded trials where the hand deviated by more than 2 cm in the wrong direction (<1% of all trials).

We used a receiver-operating characteristic (ROC) analysis to quantitatively define the presence and latency of the SLR, as described previously (Corneil et al., 2004; Pruszynski et al., 2010). We separated EMG activity based on stimulus location and SF condition. EMG activity for the same SF but opposite stimulus locations was then analyzed at every time sample (1 ms) between 100 ms before and 300 ms after stimulus presentation. For each sample we calculated the area under the ROC curve, which indicates the probability that an ideal observer could discriminate between leftward and rightward stimulus presentations based solely on the EMG activity. A value of .5 indicates chance discrimination, whereas values of 1 or 0 indicate perfectly-correct or incorrect discrimination, respectively. Discrimination threshold was set to 0.6, which corresponds approximately to the upper 95% confidence interval determined using a bootstrapping procedure (Goonetilleke et al., 2015). ROC latency was defined as the time where the ROC time series surpassed threshold and remained above it for a minimum of eight out of ten samples.

A representation of trial-by-trial EMG activity, mean EMG, and the associated time-series ROC analysis which we use to define SLR characteristics is shown for a representative participant in **Fig. 1c-e**. The SLR is visible in this example as a vertical banding indicating a change in muscle recruitment aligned to stimulus as opposed to movement onset (**Fig. 1c**), and as a transient increase or decrease in activity in mean EMG activity for leftward and rightward stimuli, respectively (**Fig. 1d;** highlighted by the grey boxes). To determine if the initial change in EMG activity was more locked to stimulus onset than movement onset, for each SF we performed a median-split of trials based on RT and conducted separate ROC time-series analyses for both ‘early RT’ (i.e., the subset of shorter RT trials) and ‘late RT’ (the subset of longer RT trials) subsets (Goonetilleke et al., 2015; Wood et al., 2015). We then calculated the slope of the line connecting the average RT and the SLR latency for the ‘early’ and ‘late’ RT sub-groups (see **Fig. 3c** for examples). Participants were classified as exhibiting a SLR if the slope of the line exceeded 67.5 degrees, which is halfway between a line parallel to the line of unity (45 degrees) and vertical (90 degrees); a slope > 67.5 degrees indicates that EMG recruitment was more locked to stimulus rather than movement onset. If a SLR was detected using the median split analysis, a subsequent time-series ROC plot was constructed using all trials to determine the SLR latency (the first of 8 of 10 consecutive points > 0.6) and the SLR magnitude (defined as the area between leftward and rightward mean EMG traces calculated from interval spanning from the SLR latency to 15 ms later).

**Figure 3.**
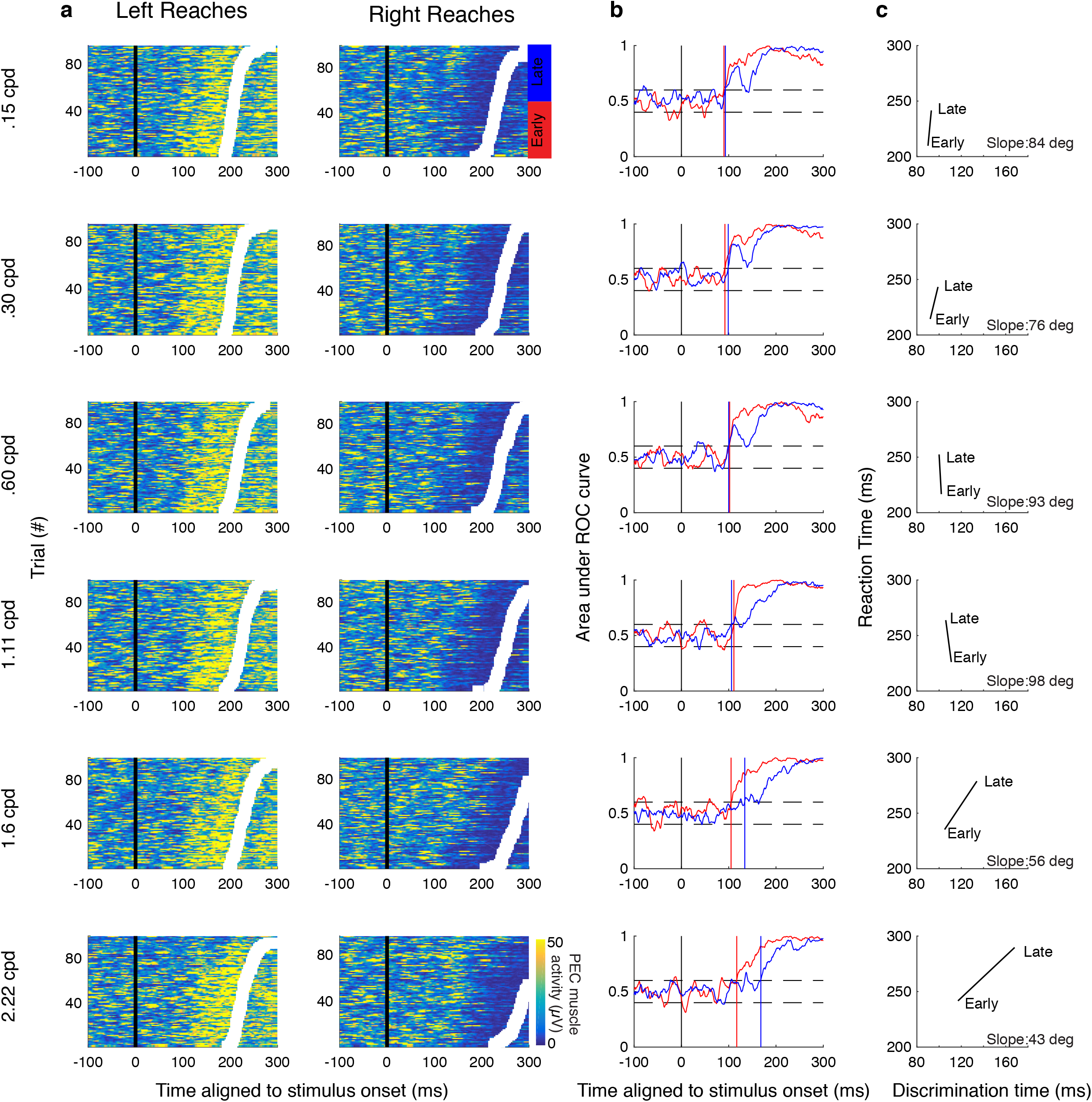
Example results from Experiment 1a. (a) Muscle recruitment from an example subject from Experiment 1a, segregated by movement direction and SF. Same format as Fig. 1c. Trials were separated into early (red) or late (blue) RT groups based on the median RT. (b) Time series ROC plot analysis from early (red) or late (blue) RT groups for data in (a). Vertical coloured solid lines depict discrimination time. (c) Plot of the average RT versus discrimination time for early and late groups. The data was deemed to exhibit an SLR if the slope of the line exceeded 67.5 deg, which indicated that the initiation of EMG recruitment was more locked to stimulus rather than movement onset.

In experiment 2, we also conducted a time-series ROC analysis to determine the latency of on-line reach corrections, based on the lateral position of the manipulandum (position along the x axis) towards leftward or rightward stimuli on jump trials. As with the analysis of EMG activity, the threshold was set to 0.6, and the latency was determined as the first of 8 of 10 consecutive points exceeding this threshold. We also defined the magnitude of the reach correction as the area between the mean trajectories for left and right reach corrections in an interval spanning from the time of discrimination to 400 ms after stimulus onset. Thus, larger magnitudes are indicative of an increased ability to move laterally towards a target.

#### Statistical analyses

Statistical analyses were performed in MATLAB (version R2016a, The MathWorks, Inc., Natick, Massachusetts, United States). Results were analyzed with paired t-tests or one-way repeated measures ANOVAs, unless otherwise stated, and post-hoc tests were Bonferroni corrected where appropriate. Chi-squared tests were used to analyze changes in SLR prevalence across SF.

## Results

### Higher spatial frequencies elicit longer reaction times

We first quantified the influence of the spatial frequency (SF) of a stimulus on reaction time (RT), across all experiments and participants. Participants performed reaches from a static hand position to stimuli placed either right or left of a start position (experiment 1a; **Fig 1a,b**), right-outward or left-outward of a start position (experiment 1b; **Fig 1a,b**), or performed on-line reach corrections of an outward reach movement to targets displaced to the left or right (experiment 2; **Fig 1a,b**). As expected based on Breitmeyer (1975), and even with the perceptual contrast-matching procedure, in all experiments we found that the SF of a stimulus influenced RTs (one-way repeated measures ANOVA of mean RT; Experiment 1a; *F*_(5,75)_ = 69.44 *p*=9.83^−27; **Fig 2a** top panel; Experiment 1b; *F*_(2,30)_ = 134.04 *p*= 1.10^−15; **Fig 2a** middle panel; Experiment 2; *F*_(2,26)_ = 99.68 *p*= 6.42^−13; **Fig 2a** bottom panel). As shown in **Fig. 2a**, RTs tended to increase for higher SFs (post-hoc analyses via paired t-tests Bonferroni-corrected for multiple comparisons; the only insignificant comparison between adjacent SFs were between 0.15 vs 0.3 cpd and 0.3 vs 0.6 cpd in experiment 1a). Importantly, RT distributions for a given SF and the next-highest SF often overlapped. In a subsequent analysis, we will exploit this overlap and mitigate the potential confound of RT on the SLR.

### Earlier SLRs are preferentially evoked by lower SFs

Next, we examined the influence of the SF of a stimulus on SLR characteristics. As shown in previous work, the SLR on limb muscles consists of a brief increase or decrease in the recruitment of muscle activity (~100 ms) (Pruszynski et al., 2010). **Fig. 1c** shows one example of a prominent SLR in one representative subject from Experiment 2, who generated leftward or rightward visually-guided corrections to a .56 cpd displaced target. In this example, the SLR appears as the prominent vertical band of EMG activity that begins ~90 ms after visual target presentation, regardless of the ensuing reaction time (see **Figs. 3** and **4** for other examples of SLRs).

**Figure 4.**
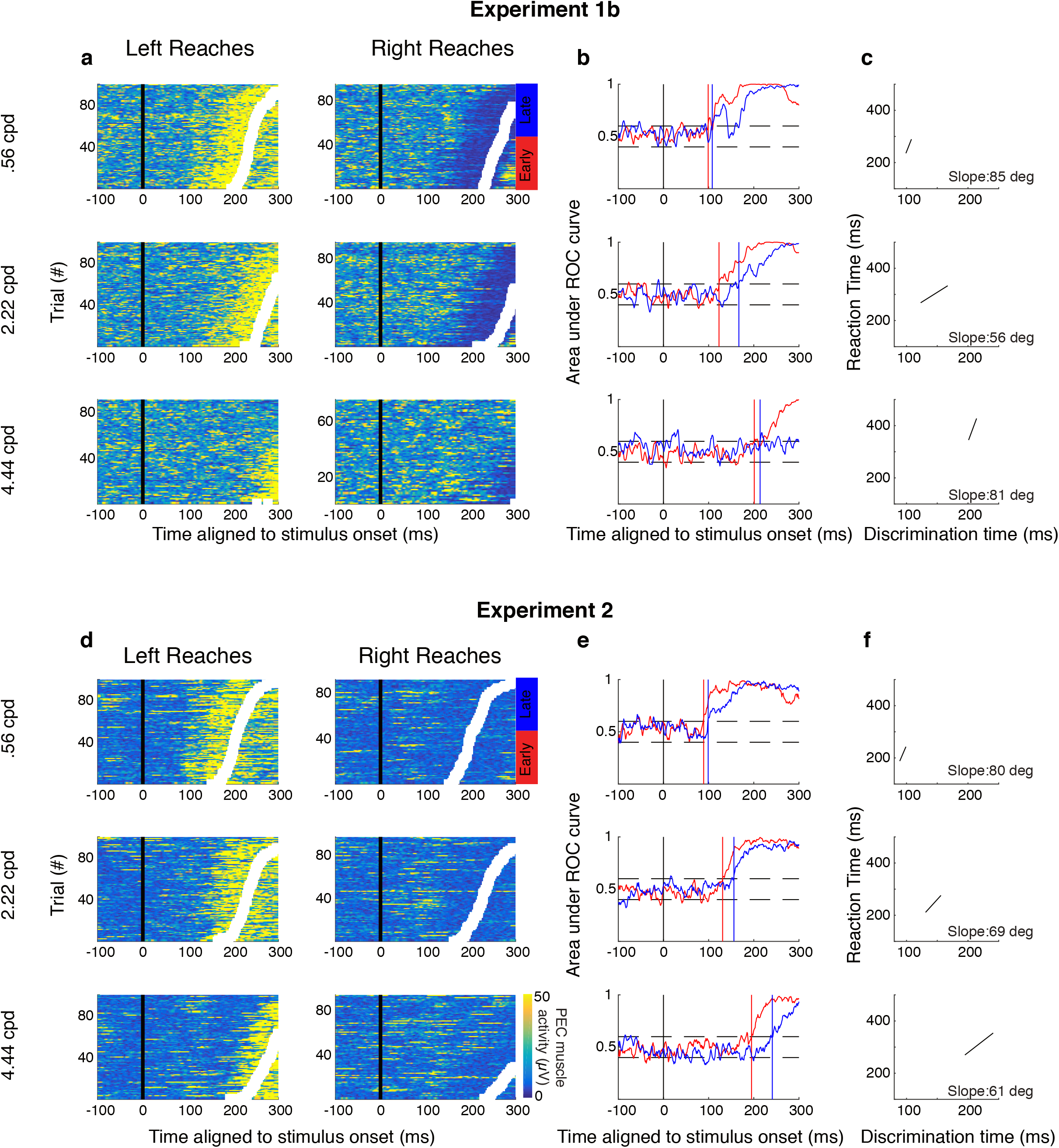
Example results from same subject in Experiment 1b (a-c) and 2 (d-f). Same format as Fig. 3.

We used a time-series ROC analysis (see Methods) to detect the presence or absence of an SLR. A participant was deemed to exhibit an SLR at a particular SF if the slope of a line connecting the reaction time plotted as a function of the discrimination time for the early and late RT groups exceeded 67.5 deg (indicating EMG activity is more locked to stimulus onset as opposed to movement onset). Using this analysis, we detected an SLR+ observation in at least one SF condition in 12 out of 16 participants in experiment 1a (**Fig. 2b** top panel; see **Fig. 3** for data from a representative subject for experiment 1a), 11 out of 16 participants in experiment 1b (**Fig. 2b** middle panel; see **Fig. 4a-c** for data from a representative subject for experiment 1b), and 12 out of 14 participants in experiment 2 (**Fig. 2b** bottom panel; see **Fig. 4d-f** for data from a representative subject for experiment 2). The data shown in **Figs. 3** and **4** also shows a qualitative pattern whereby lower SFs evoke shorter latency and more distinct SLRs when compared to SLRs evoked by higher SFs (e.g. 0.15 cpd versus 1.6 cpd rows in **Fig. 3**, .56 cpd versus 2.22 cpd **Fig. 4)**.

To quantify the influence of the SF of a stimulus on the SLR across our sample, we examined how the prevalence, latency, and magnitude of the SLR change as a function of SF. Prevalence measures the proportion of participants exhibiting an SLR at a particular SF. As shown in the top row of Fig, 5, SF influenced SLR prevalence in all experiments, with either the lowest SF (experiments 1b and 2) or second lowest SF (experiment 1a) being associated with the greatest prevalence, which was significantly different than the prevalence at the highest SF (chi-squared test; experiment 1a: *p*=.012, chi-squared= 6.35, df= 1; experiment 1b: *p*=.013, chi-squared= 6.15, df= 1; experiment 2: *p*=8.01^−5, chi-squared= 15.56, df= 1).

**Figure 5.**
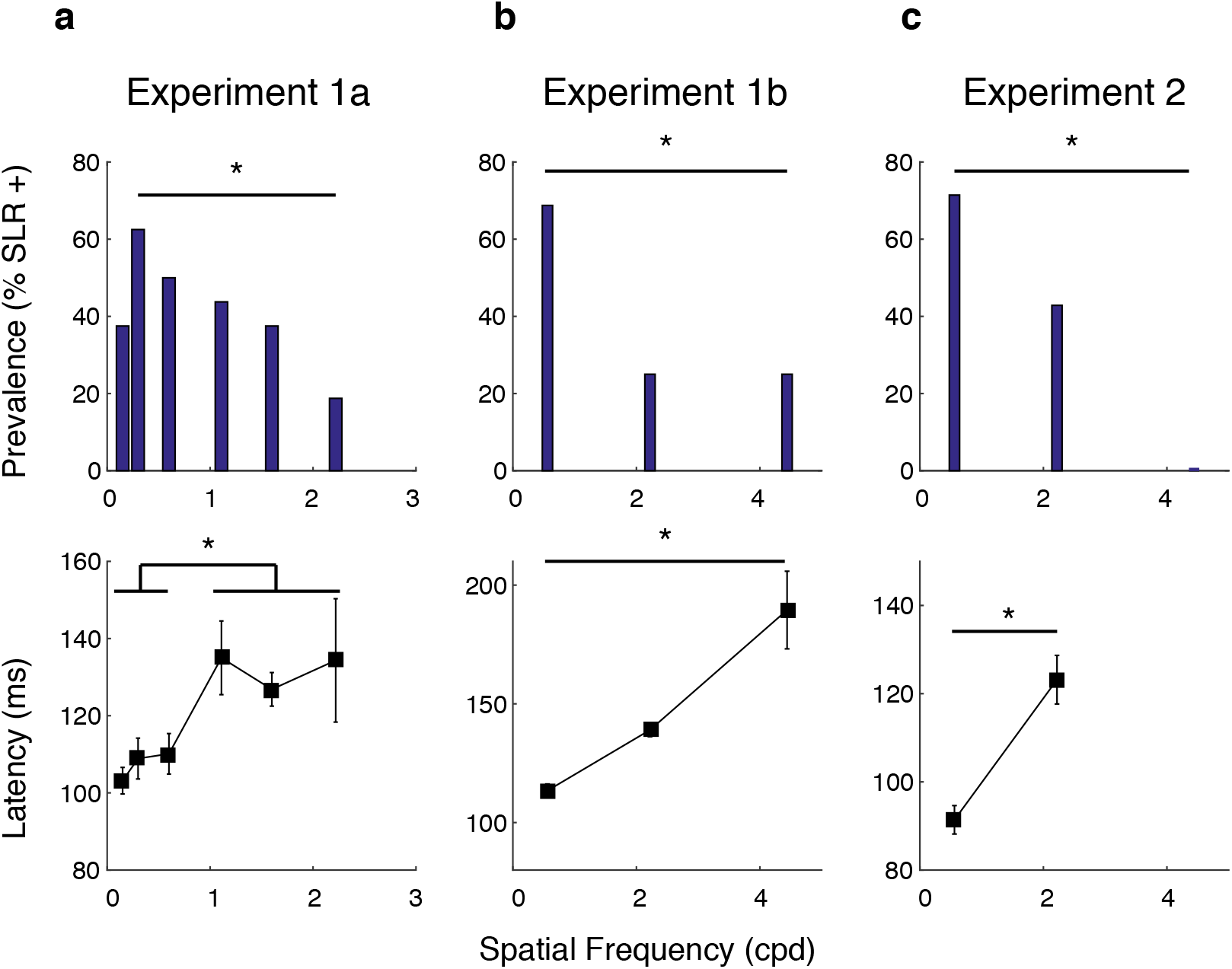
Effect of SF on latency and prevalence of the SLR, across all experiments and participants. In each column, the top row conveys SLR prevalence (the percentage of SLR+ participants across the sample) as a function of SF, and the bottom row conveys SLR latency as a function of SF. Error bars in lower row show standard error of mean. Asterisks depict differences significant at the p < 0.05 level (see Results text for details).

As shown in **Fig. 2e**, not all participants exhibited SLRs in each SF condition. This creates an unbalanced design which complicates the analysis of how the characteristics of the SLR vary as a function of SF. To analyze how SLR latency changed as a function of SF in experiment 1a, we split SLR+ observations into ‘low’ (.15, .3, .6 cpd) and ‘high’ (1.11, 1.6, 2.22 cpd) groups. For each participant, we then took the mean of any latency from an SLR+ observation for each of the ‘low’ versus ‘high’ groups, and analyzed results from participants exhibiting at least one SLR+ observation in both of the low and high conditions. For experiment 1b and 2, we analyzed results from any participant exhibiting an SLR+ observation at more than one SF. All experiments revealed an influence of stimulus SF on the latency of the SLR, with lower SFs eliciting shorter-latency SLRs (paired t-tests; Experiment 1a: t(9)= −5.37 *p*=.00045, **Fig. 5a** bottom panel; Experiment 1b: t(5)= −3.64 *p*=.015; **Fig. 5b** bottom panel; Experiment 2: t(3)= −6.13 *p*=.0087, **Fig. 5c** bottom panel).

We attempted to conduct a similar analysis on the magnitude of the SLR response, but such an analysis was complicated by the increases in SLR latency that we observed for higher SFs. As a consequence, the large volley of EMG activity associated with movement-related activity often overlapped with the SLR interval (e.g., see **Fig. 4d**, for the .56 versus 2.22 cpd condition), obscuring the quantification of EMG recruitment attributable to the SLR. SLRs tended to be easily distinguished at the lower but not higher SFs, as shown both by the vertical banding tied to stimulus onset in trial-by-trial EMG activity, and a distinct initial peak or plateau in the time-series ROC plots (see also Wood et al. 2015). In contrast, time-series ROC plots for higher SF stimuli, regardless of whether they were classified as SLR+ or SLR−, tended to lack a clear initial peak or plateau after discrimination (e.g., 2^nd^ and 3^rd^ rows of **Fig. 4e**). The lack of a clear initial peak in the time-series ROC for higher SF stimuli reinforces our concerns about whether an analysis of EMG recruitment during the SLR interval would fairly capture only activity attributable to the SLR, rather than also including movement-related activity on some subset of trials.

Overall, despite our inability to comprehensively quantify SLR magnitude in what we felt was a fair manner uncontaminated by movement-related activity, in all experiments we observed clear trends for lower SF stimuli to elicit shorter-latency and more robust SLRs.

### Latency and prevalence effects persist in results controlled for reaction time

As described in the methods, all peripheral stimuli were perceptually contrast-matched across SFs in an attempt to minimize potential perceptual confounds. Consistent with previous results, although contrast matched, higher SFs elicited longer RTs (Breitmeyer, 1975). Could the dependencies with SF noted in the previous section simply result from the generation of shorter RTs at lower SFs? To address this potential confound, we conducted an RT matching procedure in which we selected a subset of trials at different SFs with overlapping RTs. In the matching procedure, leftward and rightward reach RT data from ‘low’ SFs (the .6 cpd stimulus in experiment 1a, the .56 cpd stimulus in experiments 1b and 2) were matched to leftward and rightward reach ‘high’ SFs data (the 2.22 cpd stimulus in all experiments), respectively. In the first step of the matching procedure, we matched pairs of trials with identical RTs, without replacement. We then iteratively identified pairs of trials with RTs within +/−15 ms of each other, again without replacement, prioritizing the shortest RTs. Once this matching procedure was completed, we only analyzed datasets with more than 45 RT-matched trials.

This procedure yielded distributions with very similar RTs for leftward or rightward reaches across different SFs. The distributions of RTs matched in this way were not significantly different in 31/32 comparisons for experiment 1a, 30/32 comparisons in experiment 1b, and 26/28 comparisons in experiment 2 (*t*-test, testing at *p* < 0.05, conservatively not correcting for multiple comparisons). Of those few distributions where RTs were found to be different, mean RTs differed by less than 8 ms. We then repeated our analyses of the SLR on this subset of trials, closely-matched for RTs.

**Figure 6** shows EMG data and time-series ROC plots for a representative participant from each of the three experiments. From these examples, it is clear that the dependency of SLR with SF persists even with closely-matched RTs. For example, a clear SLR evolves after the presentation of a lower SF stimulus in experiments 1a, 1b, and 2, but not at the higher SF. Further, any features of EMG recruitment that appeared to be related to the higher SF stimulus evolved ~30 ms later than to the low SF stimulus, despite closely matched RTs (e.g., **Fig. 6g**). In these examples, the influence of SF on the earliest direction-dependent EMG activity is particularly apparent in the mean EMG and time-series ROC plots shown in the right column of **Fig. 6**, in that the divergence of EMG activity following leftward versus rightward target presentation occurs earlier when the target is composed of low than high SFs (green versus purple traces).

**Figure 6.**
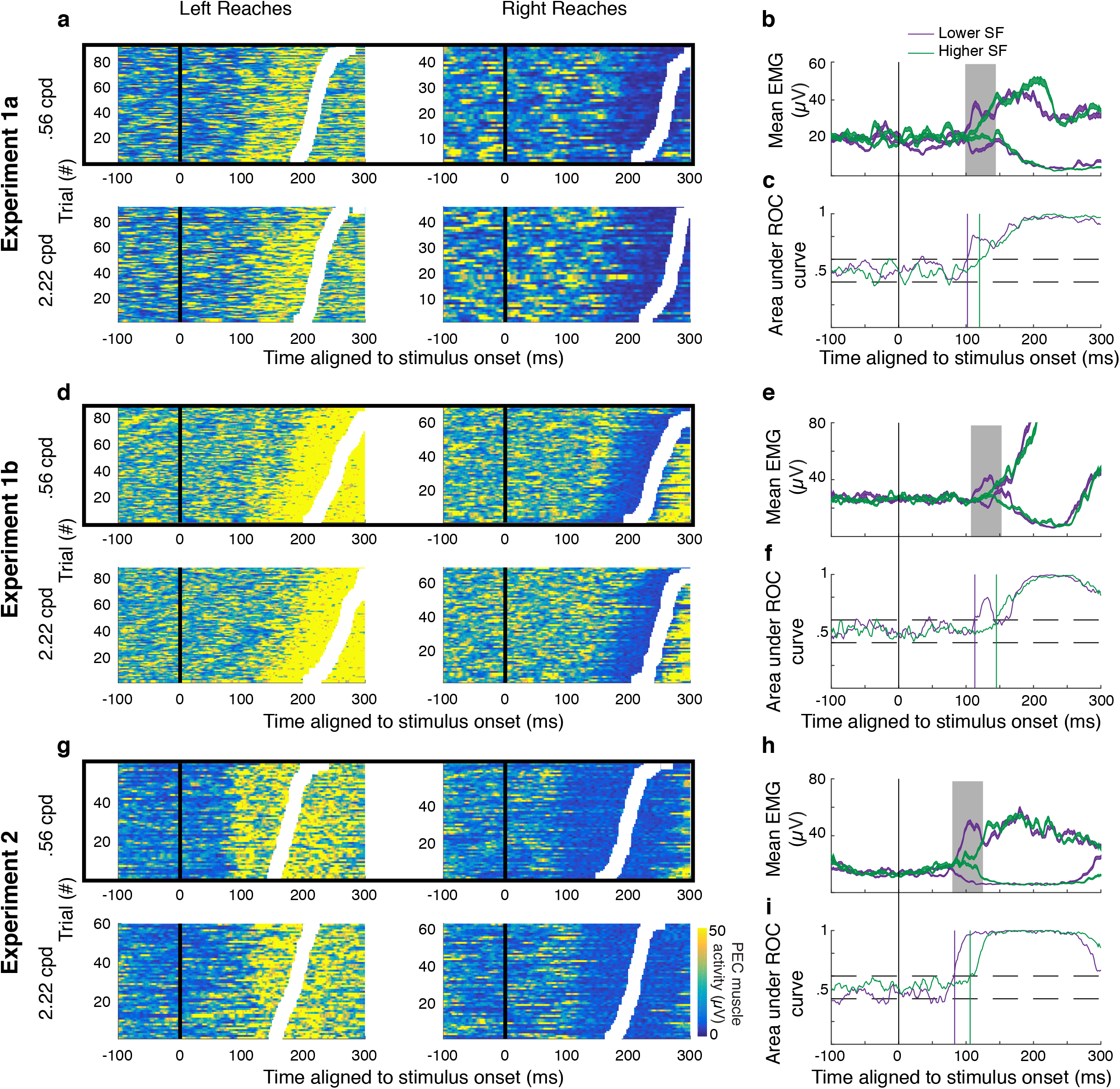
Example results from all experiments for trials matched by RTs. Trial-by-trial EMG activity (a, d, g; same format as Fig. 1c), mean EMG activity (b, e, h; same format as Fig. 1d), and time-series ROC analysis (c, f, i; same format as Fig. 1e) for data from Experiment 1a (a-c), Experiment 1b (d-f), and Experiment 2 (g-i). For each experiment, the RTs were matched for individual trials following presentation of a stimulus of a low SF (purple) or high SF (green). The solid boxes around the trial-by-trial plots depict data that exhibits an SLR, even with the reduced number of trials.

**Figure 7** shows SLR prevalence and latency across our sample for RT-matched data. From this figure it is clear that lower SFs continued to elicit more prevalent (**Fig. 7a,c,e**) and shorter latency (**Fig. 7b,d,f**) SLRs, even when RTs are closely matched. These results reached significance for SLR prevalence in experiment 2 (*p* = .018, chi-squared= 5.6, df=1), and for SLR latency in experiment 1b (t-test; t_(15)_=−3.24, *p* = .0054; **Fig. 7d**), and experiment 2 (t-test t_(8)_=−2.65, *p* = .029; **Fig. 7f**). These analyses confirm that the dependencies of SLR responses properties on the spatial frequency of a stimulus are not simply the result of short RTs to stimuli composed of lower SFs.

**Figure 7.**
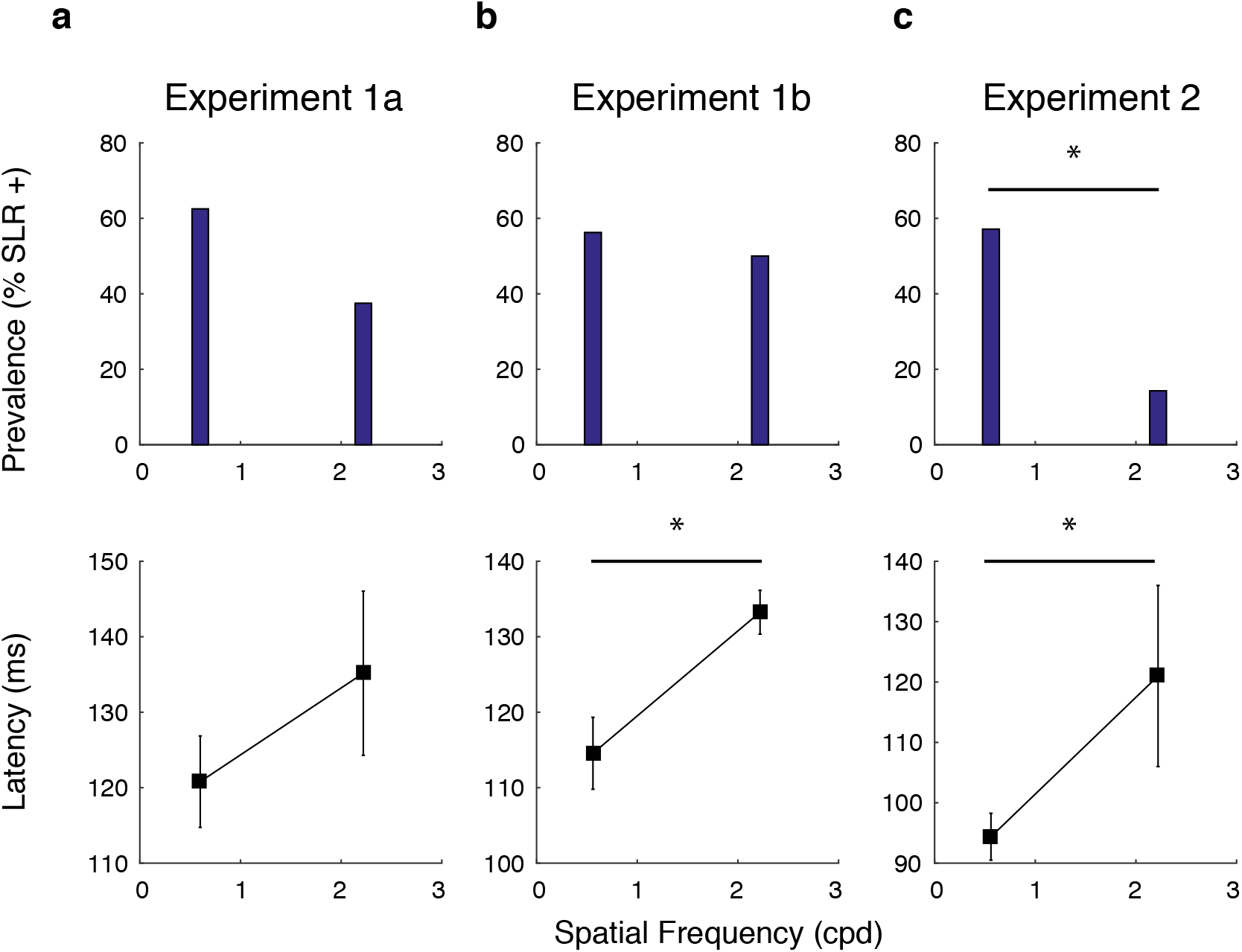
The influence of SF on SLR characteristics persisted for RT-matched trials. Same format as Fig. 5, using data matched for RT.

### SLRs, and muscle recruitment more generally, evolve earlier when the arm is already in motion

We noticed in **Figure 5** that SLR latencies were shorter when the arm was already in motion (experiment 2) versus in a stable posture (experiment 1). Recall that experiments 1b and 2 were configured so that the retinal location of peripheral targets, and the spatial position of the hand at peripheral target onset, were approximately equal (**Fig. 1a,b**). Therefore, we investigated the differences in SLR latency to the same stimulus between experiment 1b and 2. To do this, we plotted the SLR latency in experiment 2 as a function of that in experiment 1b. As illustrated in **Fig. 8a**, all but one participant response fell below the line of unity, indicating shorter SLR latencies in experiment 2 (paired t-test, t(7)= 4.88 *p*=.0018). Thus, in those few instances where an SLR was provoked to the same stimulus in both experiment 1b and 2 for the same participant, shorter SLRs were observed when the hand was already in motion (22 ± 13 ms, n = 8). Note that since these values are specifically locked to stimulus onset, this ~20 ms difference is not due to the lower RT cutoff used in Experiment 2.

**Figure 8.**
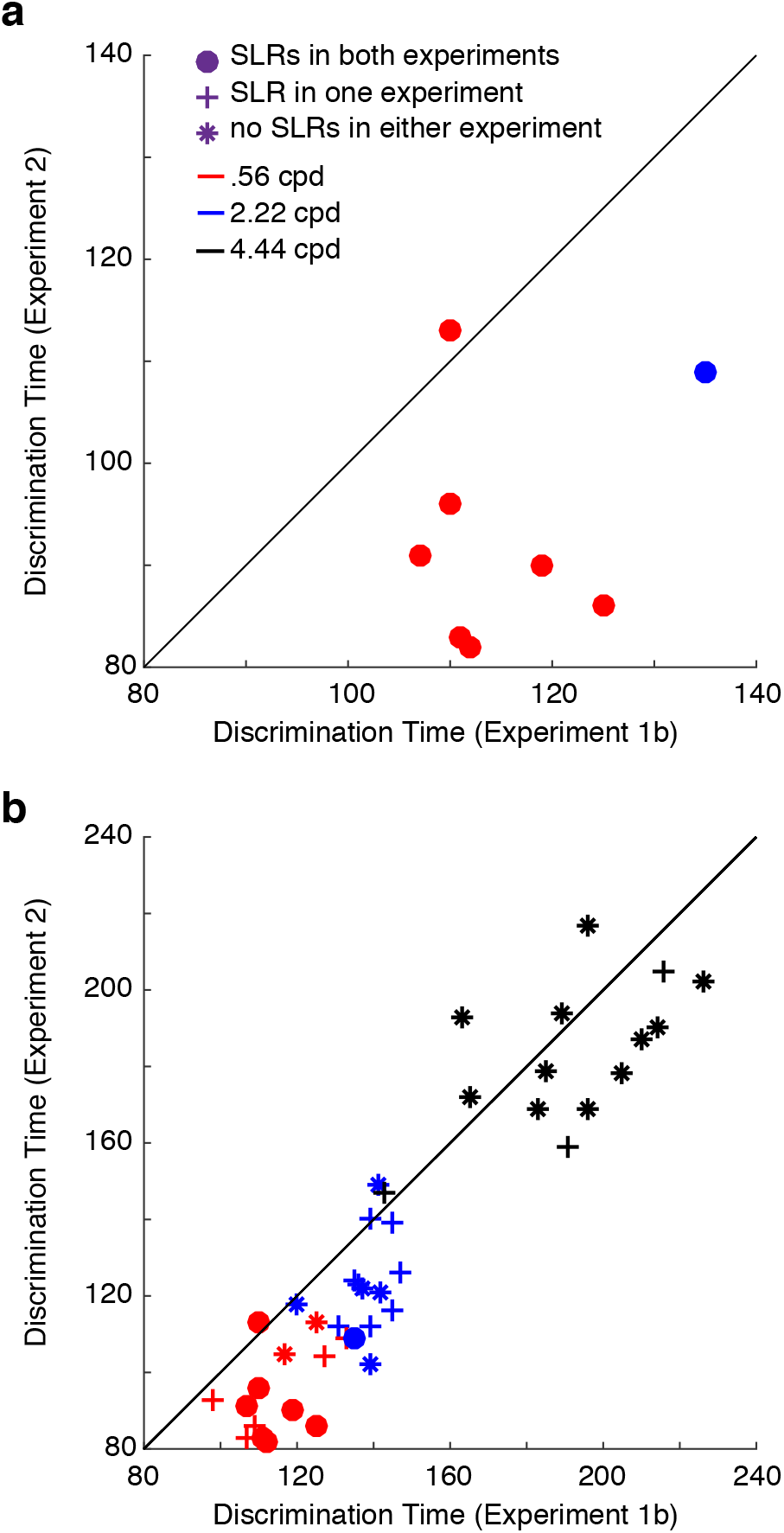
Discrimination times during on-line reach corrections (Experiment 2) plotted as a function of discrimination times for reaches initiated from a static posture (Experiment 1b). Discrimination times are derived from the time-series ROC analysis. Each data point depicts data from a unique combination of subject and stimulus SF (see colour scheme). Data from occurrences with an SLR+ observation in both Experiments 1b and Experiment 2. Data from all participants, with symbols depicting whether SLRs were observed in both experiments (dots, same data as in (a), but on a different time scale), whether an SLR was observed in only one experiment (crosses), or whether an SLR was not observed in either experiment (asterisks). Solid diagonal line shows line of unity. The clustering of data below the line of unity means that shorter discrimination times were observed in Experiment 2.

We also sought to determine the time at which EMG activity indicated the side of target presentation, regardless of whether an SLR was detected or not. As shown in **Fig. 8b**, earlier discrimination times persisted in experiment 2 across all participants and across all SFs (paired t-test, t(41)=6.25, *p* = 1.9e-7). Comparatively, we observed a smaller difference in discrimination time in those observations that lacked an SLR in both the static and dynamic experiment (13 ± 15 ms shorter in experiment 2, n = 34). Interestingly, as seen in **Fig. 8b**, SF also appeared to influence the comparative discrimination time, as the higher SFs clustered closer to the line of unity (repeated measures ANOVA *F*_(2,26)_ = 3.76 *p*= .037; **figure 8b**, red versus black differences). Thus, when the arm is in motion, information related to target presentation gets to the upper limb sooner. However, this effect is greatest in the presence of an SLR, and to low SF stimuli.

### Paralleling SLRs, on-line reach corrections start earlier for lower SF stimuli

All participants demonstrated a similar pattern of online corrections (**Fig. 9a**), wherein the lowest SF evoked the shortest latency and largest magnitude corrections. Paralleling SLR responses, the corrective latency systematically increased (one-way repeated measures ANOVA *F*_(2,26)_ = 99.78 *p*=6.34e-13, **Fig. 9b**), and magnitude systematically decreased (one-way repeated measures ANOVA *F*_(2,26)_ = 437.86 *p*=9.52e-21, **Fig. 9c**) as a function of SF.

**Figure 9.**
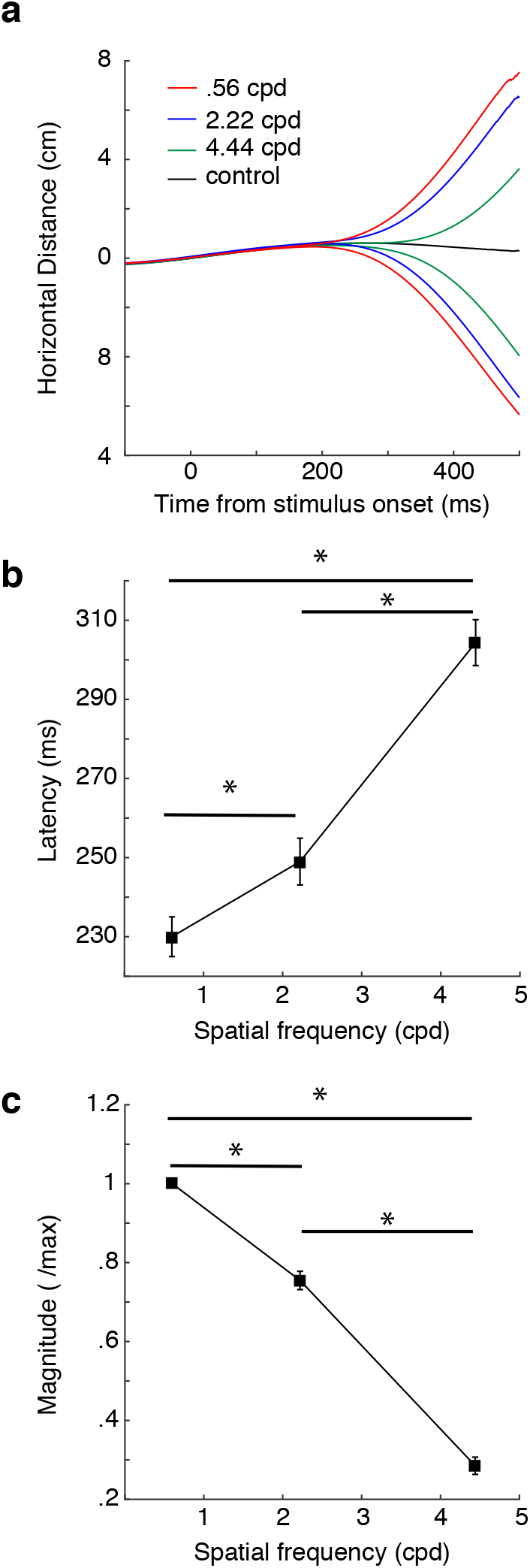
Characteristics of on-line reach corrections to different SFs in Experiment 2. Results from a sample participant, depicting the mean lateral distance for on-line reach corrections to displaced stimuli composed of different SFs (coloured lines), or to the control condition where the reach target was not displaced (black line). (b, c) Latency or magnitude (c) of on-line reach corrections across the sample, as a function of SF. The latency was derived from a time-series ROC analysis of lateral distance for left versus right displaced stimuli. The magnitude is normalized within each participant to the maximum correction. Error bars represent SEM.

Experiment 2 offers the opportunity to examine SLRs, EMG recruitment more broadly, and the parameters of on-line reach corrections in the same participants. Therefore, we collapsed across conditions to examine a range of responses. We observed that the latency and the magnitude of the online corrections were significantly correlated (Pearson’s correlation, R^2^ =.79, *p* =2.27e-15), indicating that earlier corrections related to larger lateral movement. Furthermore, as would be expected in a muscle contributing to the on-line correction, the latency of EMG discrimination was also significantly correlated with the latency of the reach correction (Pearson’s correlation, R^2^ =.77, *p* =6.06e-15), with EMG recruitment preceding kinematic changes by 128 +/− 18 ms in experiment 2. As seen in **Fig. 10**, there was also a clear three-way relationship between EMG responses, and the latency and magnitude of online reach corrections. Thus, as expected, earlier changes in EMG recruitment were associated with earlier and larger on-line reach corrections.

**Figure 10.**
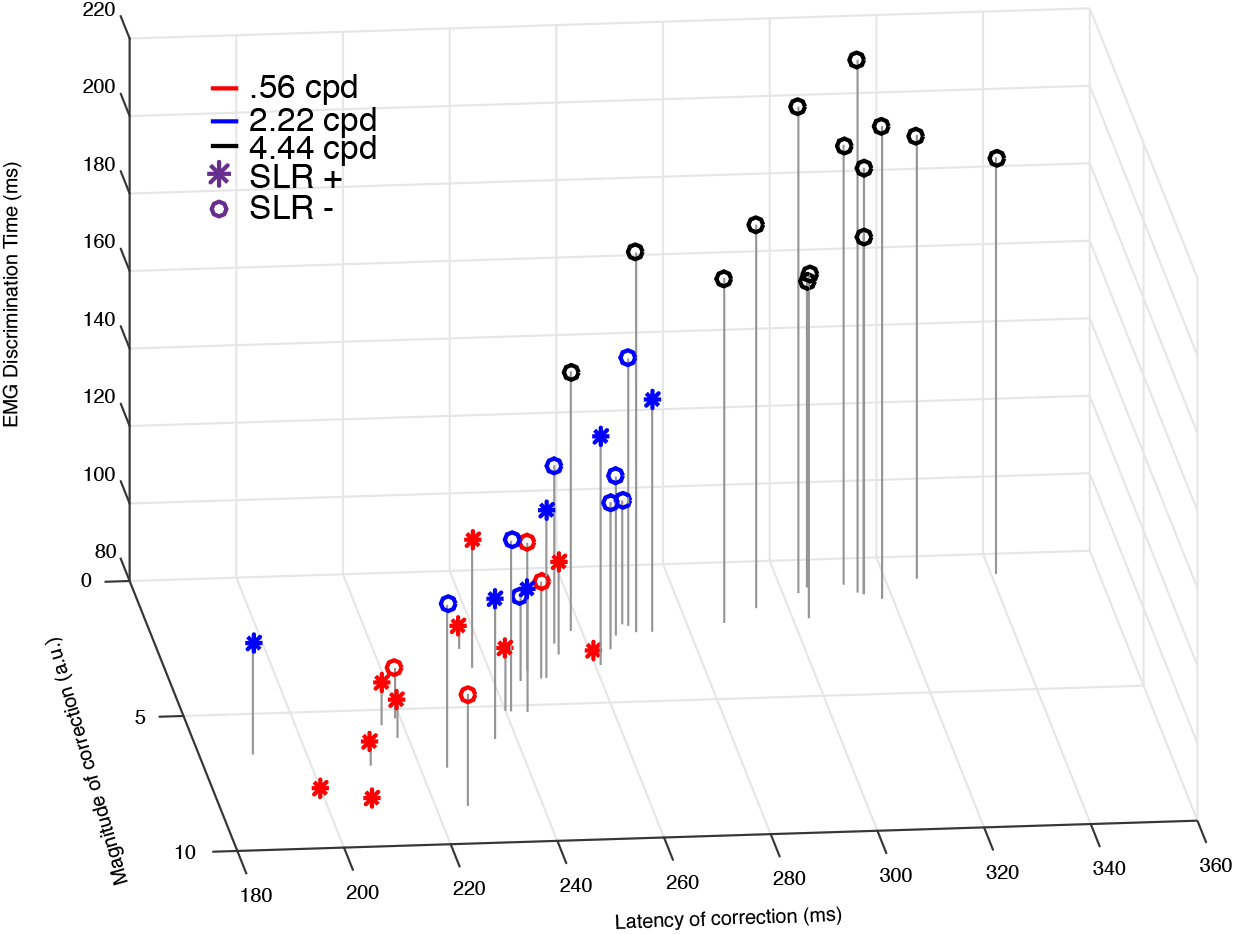
Three-way relationship between EMG discrimination, and the latency and magnitude of the on-line reach correction. Each point represents data from a single participant, colour-coded by SF, with the symbol indicating whether an SLR was observed (asterisks) or not (circles).

We also explored the relationship between the presence or absence of SLRs and online reach behavior. Firstly, the impression from **Fig. 10** is that all the data across participants falls along a smooth continuum, regardless of a response being SLR+ or not. Secondly, along this continuum, SLR+ observations tended to cluster near the most rapid and largest online corrections. To show this, we performed a *k*-means cluster analysis (with *k* = 3) and found that SLRs predominated in the clusters associated with the earliest and largest corrections (SLR prevalence of 73% (11/15), 35% (5/14), and 0% (0/13) as clusters increased in latency). Taken together, earlier EMG responses led to earlier and larger online corrections. If evoked, SLRs were associated with earlier and larger on-line corrections.

## Discussion

To interact with our visual world, we are required to transform visual information into motor commands. Here, we systematically investigated which spatial frequencies (SFs) preferentially engender rapid visuomotor responses. We analyzed stimulus-locked responses (SLRs) on upper limb muscles, as participants performed visually guided reaches towards either stationary targets from a static posture, or targets abruptly displaced while a reaching movement is in mid-flight. For all experiments, we found that low SFs elicited shorter latency and more prevalent SLRs. Paralleling SLRs, online corrections started earlier for low SFs, and consequently attained greater magnitudes. From this research, it is clear that a rapid visuomotor pathway preferentially transforms low SF visual information into the earliest phases of muscle recruitment.

### A fast visuomotor system prefers lower spatial frequencies

A variety of visual stimulus attributes, including stimulus color, intensity, and SF have also been investigated in other rapid visuomotor responses such as saccadic RTs (Bell et al., 2006; White et al., 2009; Chen et al., 2018) and ocular and manual following responses (Miles et al., 1986; Carl and Miles, 1990; Saijo, 2005; Gomi et al., 2006). These results indicate that rapid visuomotor responses rely heavily on the processing of some visual attributes, but not others. Consistent with these findings, we demonstrate that the pathway mediating rapid EMG responses and on-line corrections are preferentially sensitive to low SFs. Veerman and colleagues (2008) studied the influence of many stimulus attributes on the latency of on-line corrections and found that the shortest RT responses were generated in response to stimuli defined by contrast, orientation, or size. Importantly, although Veerman and colleagues (2008) examined reactions to a textured square, the dominant SF of the texture was not provided nor systematically varied. We further this research and show that SLRs are preferentially evoked by low SFs which may be elicited from either a static or dynamic limb position, and when present, are indicative of an increased ability to correct ongoing reach behavior towards a target.

### Plausible neural substrates for SLRs and the initial phase of on-line reach corrections

One potential structure that could serve as a sensorimotor interface for fast visuomotor responses is the superior colliculus (SC), a midbrain structure involved in coordinating the orienting reflex whereby an organism is able to rapidly realign the eyes, head, and upper body towards novel stimuli of interest (see for review (Corneil and Munoz, 2014)). There are many similarities between visual responses in the intermediate layers of the SC (SCi) and the SLR. For example, in anti-saccade or anti-reaching tasks, visual responses in the SCi (Everling et al., 1999), and the SLR on neck (Corneil et al., 2008) or limb muscles (Gu et al., 2016) are tuned to stimulus location. Visual responses present in reach-related SCi neurons also correlate well with upper limb muscle activity (Werner et al., 1997a, 1997b; Stuphorn et al., 1999), with SCi responses preceding SLRs in neck muscles by approximately ~11 ms, consistent with the efferent lag from the SCi to the neck (Rezvani and Corneil, 2008). Further, visual responses in the SCi (Li and Basso, 2008; Marino et al., 2012), SLRs on the upper limb (Wood et al., 2015), and on-line corrections (Veerman et al., 2008) all evolve at shorter latencies in response to high-contrast visual stimuli, or low spatial frequencies ((Chen et al., 2018); current results). Indeed, SLRs have been argued to resemble many aspects of express saccades (Corneil et al., 2004), when visual responses in the SCi become strong enough to drive recruitment in the motor periphery. Consistent with this perspective, express saccades have also been argued to occur at long latencies to sub-optimal stimuli (Bell et al., 2006), which has been observed in SLRs towards low contrast (Wood et al., 2015), or higher SF (**Fig. 5**) stimuli.

Our position is that SLRs, and by extension, the initial phase of online reach corrections are mediated by visual responses in the SCi. While this idea is consistent with previous ideas of how the SC may be involved in rapid visuomotor responses (Day and Brown, 2001), in general it is thought that on-line reach corrections are driven through either parietal or frontal sources (for review see (Gaveau et al., 2014; Archambault et al., 2015)). Although many of the stimulus preferences for SLRs and on-line reach corrections resemble those reported in the SCi, we are mindful that other areas in the parietal and frontal cortices implicated in on-line reach control also exhibit visually-aligned response transients (Riehle, 1991; Cisek and Kalaska, 2005), although the relationship between such transients and various stimulus features remain to be investigated. The general similarities that we are observing between SLRs and the early phase of on-line reach corrections suggests that a subcortical route involving the SCi may provide the earliest drive to the motor periphery. Subsequent, more contextually-guided portions of the online correction may rely on corticospinal inputs that incorporate processing in frontal and parietal cortices.

### Expedited visuomotor responses when the limb is already in motion

Experiments 1b and 2 were designed to compare SLRs in situations where stimulus presentation occurred when the limb was either stable or already in motion. We found that SLRs, and the earliest divergence of EMG activity, following stimulus onset evolved substantially earlier (~20-30 ms) when the limb was already in motion. This observation relates to contemporary debates about expedited RTs during movements initiated from a dynamic position (Smeets et al., 2015), and are consistent with expedited responses occurring when a postural control policy has been disengaged (Scott and Cluff, 2016). What is interesting is to consider *where* in the pathway rapidly transforming vision into actions the delay in activity aligned to stimulus onset is delayed by 20-30 ms when movements are beginning from a stable posture, and why such a delay depends on SF. It would appear that any such delay would have to be imposed downstream from the SCi, given that visual response latencies in the SCi approach the minimal afferent delay.

### Methodological considerations

Our experimental configuration limited the upper range of SFs we could present without aliasing (2.22 and 4.44 cpd in experiments1 and 2, respectively). We do not view this as a limitation, as SLRs were rarely generated in response to a 4.44 cpd stimulus. Further, any SLRs that were generated to the 4.44 cpd stimulus did so at a latency that overlapped with movement-related activity. In this vein, the potential overlap between the SLR and movement-related activity complicated analysis of SLR magnitude. Future experiments should investigate magnitude in a paradigm which separates SLR and movement related activity, where immediate reaching is not required. Interestingly, at the lower range of SFs in experiment 1a, we observed that the 0.15 cpd stimulus elicited the shortest-latency but not necessarily the most prevalent SLR, highlighting how these measures can be dissociated.

We went to some lengths to ensure the trends we observed in SLR latency and prevalence across SFs were not confounded by RT, implementing a contrast-matching procedure and reanalyzing SLRs after matching RTs. Our contrast-matching procedure also ensured that our observed effects were due to the SF of the targets, as opposed to perceived contrast. However, the on-line correction RTs we observed were longer than those reported previously (e.g., 125 ms in (Day and Lyon, 2000). We surmise that there are a few reasons for this discrepancy. Firstly, we observed single-trial examples with RTs below 200 ms (**Fig. 2a**), although we reported the mean correction latencies. Secondly, we excluded shorter-latency RTs to prevent overlap with movement-related activity. Most importantly, stimuli defined by texture are known to elicit longer-RT corrections (Veerman et al., 2008).

Finally, SLR prevalence never exceeded 75%, meaning that stimuli did not always elicit an SLR (see also Pruszynski et al., 2010; Wood et al., 2015; Gu et al., 2016). The failure to observe an SLR may be due to participant idiosyncrasies, or to the possibility that the SLR distributes to deeper layers of fatigue-resistant muscle (Pruszynski et al., 2010). As mentioned above, Gabor patches may also be a suboptimal stimulus for the SLR. Recall also that our classification criteria relied on a median-split analysis. This is a conservative approach that requires a substantial number of trials and considerable variance in RTs.

### Conclusions

The work presented here builds on a stream of findings that emphasize the importance of stimulus attributes on visually-evoked responses in premotor areas and the motor periphery, showing that the earliest component of a rapid visuomotor response is preferentially elicited by low SF stimuli. The data from experiment 2 are the first to directly link the phenomenon of SLRs to on-line reach corrections in the same task and in the same participants. This link will help further characterizations of the substrates underlying rapid visuomotor responses, as the SLRs can be readily elicited on a trial-by-trial basis while participants are in a stable posture. Detailing the stimulus attributes which best elicit rapid visuomotor responses will help identify plausible underlying substrates and will aid the study of such responses across the lifespan and in clinical populations.

## Acknowledgements

The authors thank Dr. Andrew Pruszynski for the use of his exoskeleton, and Rodrigo Maeda for assistance in the initial setup. This work was supported by a Discovery Grant from the Natural Sciences and Engineering Research Council (NSERC; RGPIN 311680) and an operating grant from the Canadian Institutes of Health Research (MOP-93796). RAK was supported by an Ontario Graduate Scholarship. CG was supported by an NSERC Canada Graduate Doctoral Scholarship.

